# Dopamine receptor sensitivity and Pavlovian conditioned approach

**DOI:** 10.1101/2025.09.26.678801

**Authors:** Nana K. Amissah, Jordan A. Tripi, Christopher P. King, Paul J. Meyer

**Affiliations:** Psychology, University at Buffalo, Buffalo, NY

**Keywords:** Dopamine, Receptor sensitivity, Cue-reactivity, Ultrasonic vocalization

## Abstract

Understanding the determinants of individual differences in cue-reactivity and drug sensitivity is critical to identifying neurobiological mechanisms underlying vulnerability to addiction. In this study, we examined the relationship between dopamine D1 and D2 receptor sensitivity and the attribution of incentive salience to reward cues and sensitivity to cocaine. Male Sprague Dawley rats were classified as having high or low sensitivity to the D2 receptor agonist quinpirole, and a subset was tested with the D1 receptor agonist SKF 82958. Cue-reactivity was assessed using a Pavlovian conditioned approach (PavCA) task, which distinguishes between sign-tracking (approach to a cue that predicts reward) and goal-tracking (approach to the site of reward delivery). Cocaine sensitivity was measured by locomotor activity and 50-kHz ultrasonic vocalizations (USVs), a putative measure of appetitive states. High D2 responders exhibited more sign-tracking and greater cocaine-induced USVs than low responders despite no difference in cocaine-induced locomotion. Sign-trackers also showed greater locomotor sensitivity to D1 receptor stimulation than goal-trackers and produced more cocaine-induced USVs. Rats with high sensitivity to both D1 and D2 receptor stimulation showed the strongest sign-tracking behavior and affective response to cocaine. These findings suggest that dopamine receptor sensitivity is associated with the propensity to attribute incentive salience to reward cues and potentially the appetitive effects of cocaine. This dopaminergic phenotype may reflect a mechanism contributing to both individual differences in cue-reactivity and drug responsiveness.

## Introduction

Decades of addiction research have been focused on understanding the transition from casual to uncontrollable drug use through the lens of drug-induced changes in neural signaling and long-term neuro-adaptations following chronic use (Deroche-Gamonet & Piazza, 2014; Hogarth et al., 2013; Koob, 2013; Meyer et al., 2015; Ostlund & Balleine, 2008; Piazza & Deroche-Gamonet, 2013; M. J. F. Robinson et al., 2013). However, not all individuals who are exposed to drugs of abuse develop substance use disorder. Epidemiological studies indicate that, depending on the drug, about 17% of individuals who have tried illicit drugs, such as cocaine, go on to fit the criteria for this disorder within 10 years (Anthony et al., 1997; SAMHSA, 2025). This highlights the inter-individual differences in risk and resilience, despite drug history, and further, emphasizes the importance of understanding the inherent genetic, neural, and behavioral factors that are predictive of continued use following initial exposure.

Clinical and preclinical studies have examined biological and behavioral factors that predispose subjects to exhibit addiction-like behaviors. For example, a subset of individuals display similarities within the mesolimbic dopamine system (Blum et al., 1990; Volkow et al., 2002a), which is important because dopamine neurotransmission plays a major role in the reinforcing effects of drugs of abuse and reward-related stimuli (Dang et al., 2018; Volkow et al., 2006). Variability in dopaminergic function, particularly at the receptor level, has been associated with individual differences in initial response to drugs and drug cues as well as a predictive factor for addiction-related phenotypes (Baik, 2013; Huys et al., 2014; Merritt & Bachtell, 2013a; Volkow et al., 1999). For example, Tournier et al. (2013) determined that innately low dopamine D2 receptor availability was related to high novelty-seeking and also to increased behavioral sensitization to the stimulant amphetamine. Additional studies have confirmed that low baseline D2 receptor availability is correlated with increased rates of cocaine self-administration in primate and rodent models (Dalley et al., 2007; Nader et al., 2006), and also high ratings of initial drug-liking in humans (Volkow et al., 2002b) that in turn is predictive of abuse potential (Davidson et al., 1993).

Further compelling evidence for the relationship between dopamine D2 receptor function, drug reinforcement, and addiction liability has been demonstrated by Merritt and Bachtell (2013), in which sensitivity to the D2 receptor agonist quinpirole was predictive of cocaine-induced behaviors. In this study, individuals characterized as high responders following a quinpirole dose-response test displayed increased time spent in a cocaine-paired context in a conditioned place preference (CPP) paradigm and increased cocaine self-administration under an FR5 schedule of reinforcement, compared to low quinpirole-responding rats. These results support the notion that dopamine D2 receptor sensitivity is related to the response to cocaine and cocaine-paired cues. This also suggests that D2 receptors affect 1) the attribution of incentive salience to reward-related cues in that individuals with high D2 sensitivity are more likely to assign motivational value to drug cues (Tunstall & Kearns, 2015), and 2) the initial subjective effect of the drug in that high D2 sensitivity is related to a heightened positive experience during initial exposure.

These processes may be acting separately or in tandem to promote the cocaine-seeking and taking observed in Merritt and Bachtell (2013). For example, the ability of a reward-associated cue to acquire its own motivational value and promote reward cue seeking is dependent on dopamine receptor function (Danna & Elmer, 2010; Flagel et al., 2011; Fraser et al., 2016). Further, there are substantial differences among individuals in terms of whether cues acquire this motivational value, which can be measured using a Pavlovian conditioned approach (PavCA) procedure. During this procedure, presentations of a lever-cue are followed by a food reward for 25 trials a day over 5 days, during which a subset of subjects come to approach and interact with lever-cue (sign-tracking), while others approach the food-delivery location (goal-tracking) (Boakes, 1977; Flagel & Robinson, 2017; Hearst & Jenkins, 1974; Meyer et al., 2012a; Robinson et al., 2014; Robinson & Flagel, 2009). Studies using this model of cue-reactivity have determined that the sign-tracking phenotype is related to enhanced drug-directed and drug cue-directed behavior, and thus these subjects may be more vulnerable to the development and maintenance of addiction-related behaviors (Meyer, 2012b; Robinson et al., 2014; Saunders & Robinson, 2010; Versaggi et al., 2016; Yager & Robinson, 2013). Particularly relevant to the current study is the finding that sign-tracking is related to the response to cocaine cues in a Pavlovian paradigm (Meyer, 2012b). Further, dopamine D2 receptor function is selectively involved in the acquisition of sign-tracking (Roughley & Killcross, 2019) and in situ hybridization studies have indicated that, compared to rats that primarily exhibit goal-tracking, rats exhibiting sign-tracking behavior display lower dopamine D2 receptor mRNA expression, but a greater proportion of striatal D2 receptors in the high affinity state (Flagel et al., 2007, 2010).

Taken together, these results suggest that differences in D2 receptor sensitivity may underlie differences in reactivity to both food- and cocaine-cues. Whereas D2 receptor sensitivity is typically measured by alterations in locomotion, cocaine produces other unconditioned effects such as the production of frequency-modulated 50 kHz ultrasonic vocalizations (Browning et al., 2011; Ma et al., 2010; Meyer, 2012b), a proposed ‘self-report’ of motivation and positive affective states in rodents (Brudzynski, 2013; Knutson et al., 2002; Mahler et al., 2013). Drug-induced USVs require dopamine neurotransmission via dopamine receptors and have been found to be related to the rate of acquisition of cocaine self-administration (Browning et al., 2011; Wright et al., 2013) as well as cocaine conditioned place preference (Meyer et al., 2012b). Further, the sign-tracking phenotype is related to increased acute and chronic cocaine-induced USVs, and it may be that these behaviors rely on overlapping neurobiological mechanisms (Meyer et al., 2012b; Meyer & Tripi, 2018; Tripi et al., 2017).

In order to determine whether dopamine D2 receptor sensitivity is related to reward cue-reactivity and the initial subjective response to cocaine, we characterized Sprague-Dawley rats as high or low responders to the D2 receptor agonist quinpirole and then tested for their tendency to attribute incentive salience to a reward-associated cue in Pavlovian conditioned approach and their response to cocaine as measured by locomotor behavior and 50 kHz USVs. Further, to test whether dopamine D1 receptor sensitivity is similarly related to these behaviors (Fraser et al., 2016; Macpherson & Hikida, 2018), we also tested a subset of subjects in their locomotor response to the dopamine D1 receptor agonist SKF 82958.

## Methods and Materials

### Subjects and housing

Male Sprague Dawley rats (*n* = 48; 250 - 275g) were purchased from Envigo (Indianapolis, IN). Upon arrival, subjects were pair-housed in a temperature- and humidity-controlled room on a reverse 12 h light-dark cycle (lights off at 0730) and handled daily for one week. All testing occurred during the dark phase of this cycle. In their home cages, food and water were available *ad libitum* throughout the study. All experimental procedures followed the guidelines of laboratory animal care specified and approved by the Institutional Animal Care and Use Committee (IACUC) at the University at Buffalo.

### Drugs

All drugs including selective D2-like receptor agonist quinpirole HCl, selective D1 receptor agonist SKF 82958 HBr (Tocris, Minneapolis, MN) and cocaine HCl (Nat. Inst. of Drug Abuse, Bethesda, MD) were dissolved in 0.9% saline for preparation of treatment doses prior to testing.

### Apparatus

#### Locomotor chambers

Locomotor chambers were constructed with black acrylic walls (47.5 cm length x 15.5 cm width x 30 cm height) and smooth matte black flooring. Positioned directly above each chamber were infrared video cameras hooked up to a 16-channel DVR (Swann Communications, Inc., Santa Fe Springs, CA). Locomotion was captured by the cameras and analyzed by Topscan video tracking software (Clever Sys., Inc., Reston, VA) (Flagel et al., 2007; King et al., 2017; Meyeret al., 2012b). This same apparatus was also used for all locomotor testing.

#### Pavlovian conditioning chambers

Med-Associates conditioning chambers (20.5×24.1 cm floor area, 29.2 cm high; St. Albans, VT) were equipped with LED-illuminated retractable levers located on either the left or right side (counterbalanced across rats) of a central food cup (3 cm above a stainless-steel grid floor). An illuminated red house light was located on the chamber wall opposite the food cup. Banana-flavored food pellets (45 mg, BioServ, #F0059, Frenchtown, NJ) were delivered into the cup by an automatic feeder. The food cup was equipped with an infrared photo-beam to detect head entries.

### Procedures

#### Quinpirole serial dose-response test

This procedure was adapted from Merritt and Bachtell (2013). Prior to each testing session, rats were removed from their home cages, weighed, and then transported to the testing room. All subjects were first habituated to the locomotor chambers for 1-hour post saline injection (0.9%, *s*.*c*.). On the following day, subjects underwent a 4 hour within-session dose response test in which locomotion was measured for 1 hour in response to saline (*s*.*c*), and then for 1 hour to each quinpirole dose administered in ascending order (0.1, 0.3, 1.0 mg/kg, *s*.*c*.). Rats’ quinpirole sensitivity was calculated using area-under-the-curve of total quinpirole-induced locomotion. A median split of this measure allowed for the classification of rats as high and low responders (D2 Hi and D2 Lo, respectively.

#### Pavlovian Conditioned Approach

One week following the quinpirole sensitivity test, rats were given ∼50 food pellets in their home cages on the two days prior to the start of Pavlovian Conditioned Approach (PavCA) (Flagel et al., 2007; Meyer et al., 2012a). Upon first exposure to the PavCA apparatus, rats underwent a single day of food cup training in which 25 food pellets were dispensed into the food cup on a variable time 30 (1-60 s) schedule. The lever was retracted, and the house light remained on for this session. Beginning on the subsequent day, rats were trained in five sessions of PavCA (one session/day). Each session consisted of 25 trials in which an 8s illuminated lever presentation (CS) immediately proceeded with the delivery of a food pellet (US). Trials were separated according to a variable time 90 (30-150 s) schedule. It is important to note that the delivery of a food pellet was never contingent on the subject’s behavior. Both lever contacts and food cup entries were measured with sessions lasting 37.5 min on average.

In addition to raw measures of lever contacts and food cup entries, tendency to sign- or goal-track can also be captured using the PavCA Index, as described by Meyer et al., 2012a. Briefly, the PavCA index, ranging from -1.0 to 1.0, is calculated from the average of three factors per session; *response bias* [(lever presses minus CS food cup entries) / (lever presses plus food cup entries)], *approach probability difference* [(# of trials with at least one lever press minus # of trials with at least one food cup entry during the CS) / 25 trials], and *latency difference* [(latency to enter the food cup during the CS minus latency to lever press) / 8]. The average of this PavCA index score from the last 2 sessions (day 4 and 5) is used to determine an overall index, thus characterizing rats as sign-trackers (ST; PavCA Index ranging from 0.5 to 1.0) or goal-trackers (GTs; -0.5 to -1.0). Rats not meeting these criteria are characterized as intermediates (IN; PavCA Index -0.49 to 0.49).

#### Cocaine-treatment

Five days following PavCA testing, rats were removed from home cages, weighed and then transported to the testing room for two consecutive days of testing. On day 1, rats received an injection of saline immediately prior to being placed in the locomotor chambers for 30 min. On day 2, all rats were given a cocaine injection (10 mg/kg, *i*.*p*.) before being placed in the chamber. In addition to locomotion, USV production was recorded using condenser microphones with a flat frequency range up to 250 kHz at 8-bit resolution (Model CM16/CMPA, Avisoft Bioacoustics, Berlin, Germany), which were placed 8 cm above each chamber and connected to an UltraSoundGate 416H recorder with four balanced analog inputs (Avisoft Bioacoustics, Berlin, Germany). USV recordings were collected and converted to spectrograms to be analyzed by an experimenter in Adobe Audition (Adobe Systems Inc, San Jose, CA).

#### SKF 82958 serial dose-response test

Three days following the cocaine test, rats classified as sign- and goal-trackers (n = 28) underwent a final test to determine sensitivity to the D1-like receptor agonist SKF 82958. For this, rats were again habituated to the locomotor chambers under saline conditions in one 50 min session. On the following day, subjects underwent a 2.5 hour within-session dose response test in which locomotion was measured for 50 min in response to saline (*i*.*p*.), and then for 50 min to each SKF 82958 dose administered in ascending order (0.1, 1.0 mg/kg, *i*.*p*.). Doses were selected based on ability to stimulate locomotor activity within the first hour following administration (Desai et al., 2005; Nergårdh et al., 2005).

### Statistics

For both the D1 and D2 agonist dose-response tests, differences in locomotion were measured using repeated-measures analysis of variance (RM ANOVA) with *PavCA Phenotype* (ST, GT) as the between groups variable, and *Dose* (4 quinpirole doses or 3 SKF 82958 doses) and *Time* (12 5-min periods for quinpirole, 10 5-min periods for SKF 82958) as the within-group variables.

Bonferroni-corrected post-hoc tests were used to determine significant differences between groups at specific doses. For PavCA measures, approach behaviors and index scores were assessed using RM ANOVA, using *PavCA Phenotype* (ST, GT) or *D2 Sensitivity* (Hi, Lo) as the between-groups variable, and Day (1-5) as the within-groups variable. Similarly, cocaine-induced behaviors were measured using the same test and between-groups variables, with Drug (saline, cocaine) as the within-groups variable.

Finally, RM ANOVA was used when considering the contribution of D1 and D2 agonist sensitivity in PavCA and cocaine-induced behaviors. For these analyses, D1 Sensitivity (Hi, Lo) and D2 Sensitivity (Hi, Lo) were used as between-group variables, and Day (1-5) or Drug (saline, cocaine) were used as within-group variables. One subject was determined to be an outlier (z-score = 4.18) based on locomotor response to quinpirole and thus was dropped from all analyses. Statistical comparisons were considered significant at p > 0.05.

## Results

### Locomotor Response to the D2 receptor agonist quinpirole, calculation of area under the curve

Consistent with Merritt and Bachtell (2013), rats displayed variation in their locomotor response to the 3 doses of quinpirole. For each subject, total quinpirole-induced locomotor activity for the two locomotor activating doses (0.3 mg/kg and 1.0 mg/kg) was assessed based on area under the curve. A median split of this value for each subject allowed for the classification of rats as either high D2 responders (Fig. 1; D2 Hi; n = 23) or low D2 (D2 Lo; n = 24) responders. As expected, individuals characterized as D2 Hi displayed an increased locomotor response to quinpirole compared to individuals characterized as D2 Lo. All main effects and interactions were significant, including the [*D2 sensitivity* x *Dose x Time* interaction: *F* (22, 990) = 7.46, p < 0.001]. However, unlike Merritt and Bachtell (2013), we found a significant difference in the response to a saline injection (not shown, F (1, 45) = 73.45, p < 0.001).

**Fig. 1.**
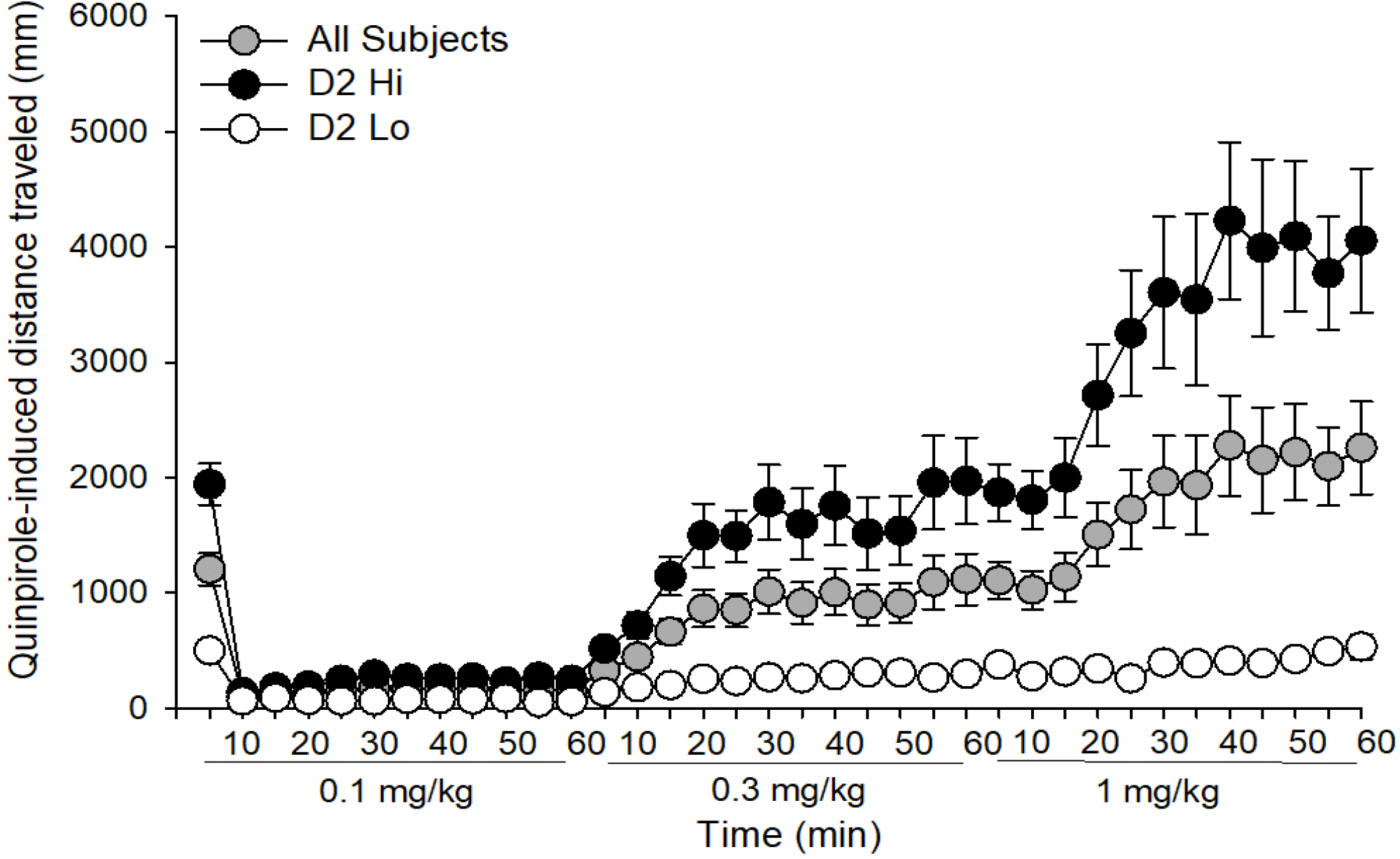
Locomotor response to quinpirole was used to characterize rats as sensitive (D2 Hi) or insensitive (D2 Lo) to a D2 receptor agonist based on a median split of area-under the curve as described in *Methods and materials* (median score = 16305.45). Data are represented as the mean (+/-SEM).

### Pavlovian Conditioned Approach Behaviors in D2 Hi and Lo rats

Lever contacts and food cup entries were measured across 5 days of Pavlovian conditioned approach (PavCA; Fig. 2). While D2 Hi and D2 Lo rats both increased lever contacts across testing days [main effect of *Day*: *F* (4, 180) = 29.38, p < 0.001], D2 Hi rats contacted the lever more often (Fig. 2A) [main effect of *D2 Sensitivity*: *F* (1, 45) = 6.51, p < 0.05]. There was no *D2 Sensitivity x Time* interaction. In contrast, D2 sensitivity did not predict food-cup entries across days (Fig. 2B). To further assess the differences in PavCA conditioned behaviors between D2 sensitivity groups, PavCA index scores were calculated for individuals on each of the five days using the equation described in the methods. There were changes in index scores across days [main effect of *Day*: *F* (4, 180) = 5.94, p < 0.001] and PavCA index scores were greater in D2 Hi rats compared to D2 Lo rats [main effect of *D2 Sensitivity*: *F* (1, 45) = 7.33, p < 0.01)], indicating a bias towards sign-tracking behavior in D2 Hi rats (Fig. 2C). There were no *D2 Sensitivity x Day* interactions.

**Fig. 2.**
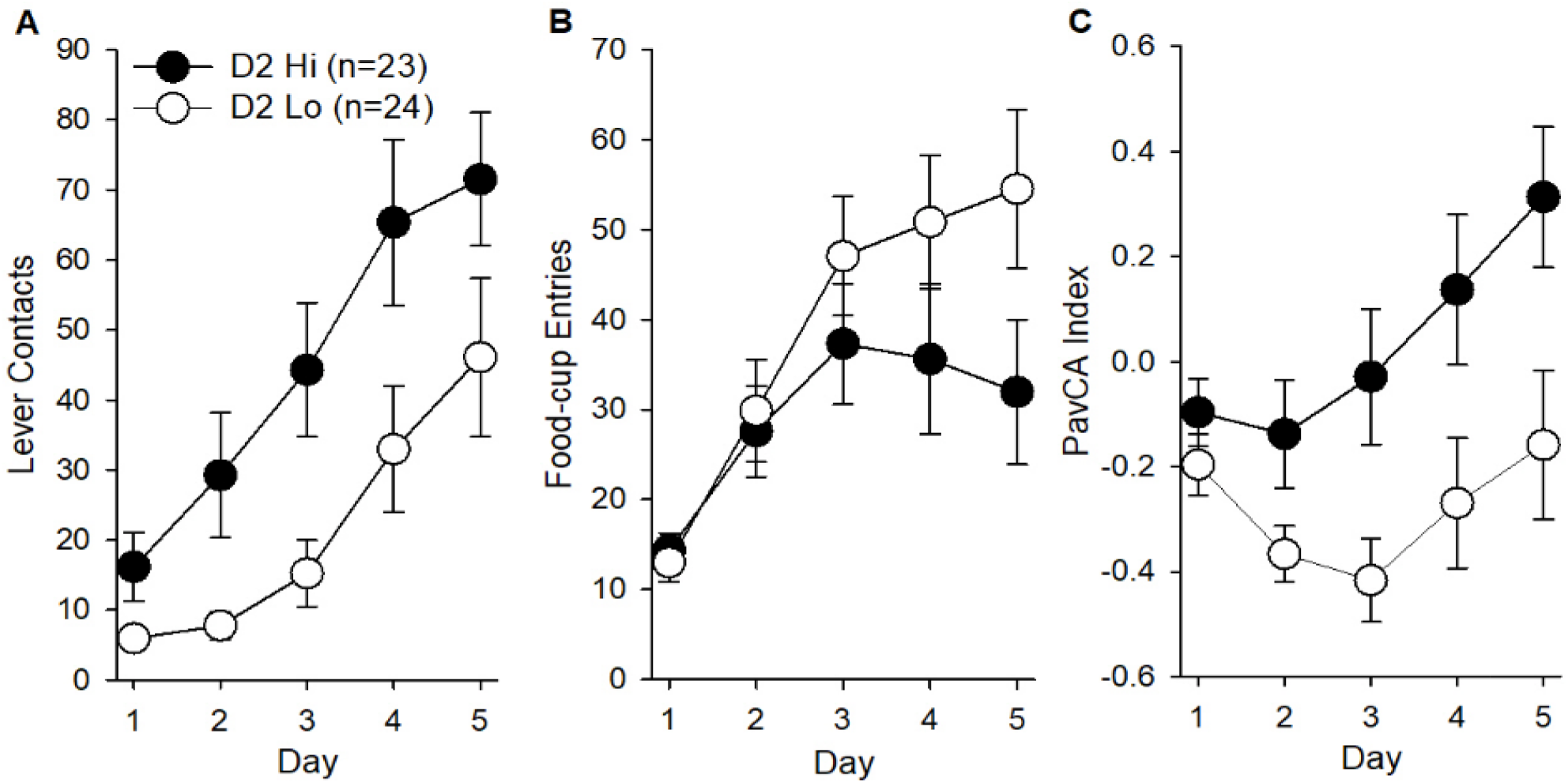
Pavlovian conditioned approach measures in D2 Hi (n=23) and D2 Lo (n=24) rats including mean number of lever contacts *F* (4, 180) = 29.38, p < 0.001 (A) and food-cup entries (B) are shown. Mean PavCA index scores *F* (1, 45) = 7.33, p < 0.01 (C) were determined using the formula presented in *Methods and materials*. Data are represented as the mean (+/-SEM).

### Cocaine-induced locomotion and USV production in D2 Hi and Lo rats

Following PavCA testing, locomotion and production of 50 kHz ultrasonic vocalizations (USVs) were measured in subjects following saline and cocaine treatment on two consecutive days. The locomotor data for four rats were removed from the analysis because of technical problems during the saline test day. Cocaine increased locomotion and USV production compared to saline treatment [main effects of *Drug*: *F* (1,41) = 12.12, p < 0.01; *F* (1,45) = 35.1; p < 0.001, respectively]. While there were no group differences in cocaine-induced locomotion (Fig. 3A), there was a trend towards a significant D2 sensitivity x Drug interaction for USVs (p =0.053) with D2 Hi rats tending to produce more cocaine-induced USVs compared to D2 Lo rats (Fig 3B).

**Fig. 3.**
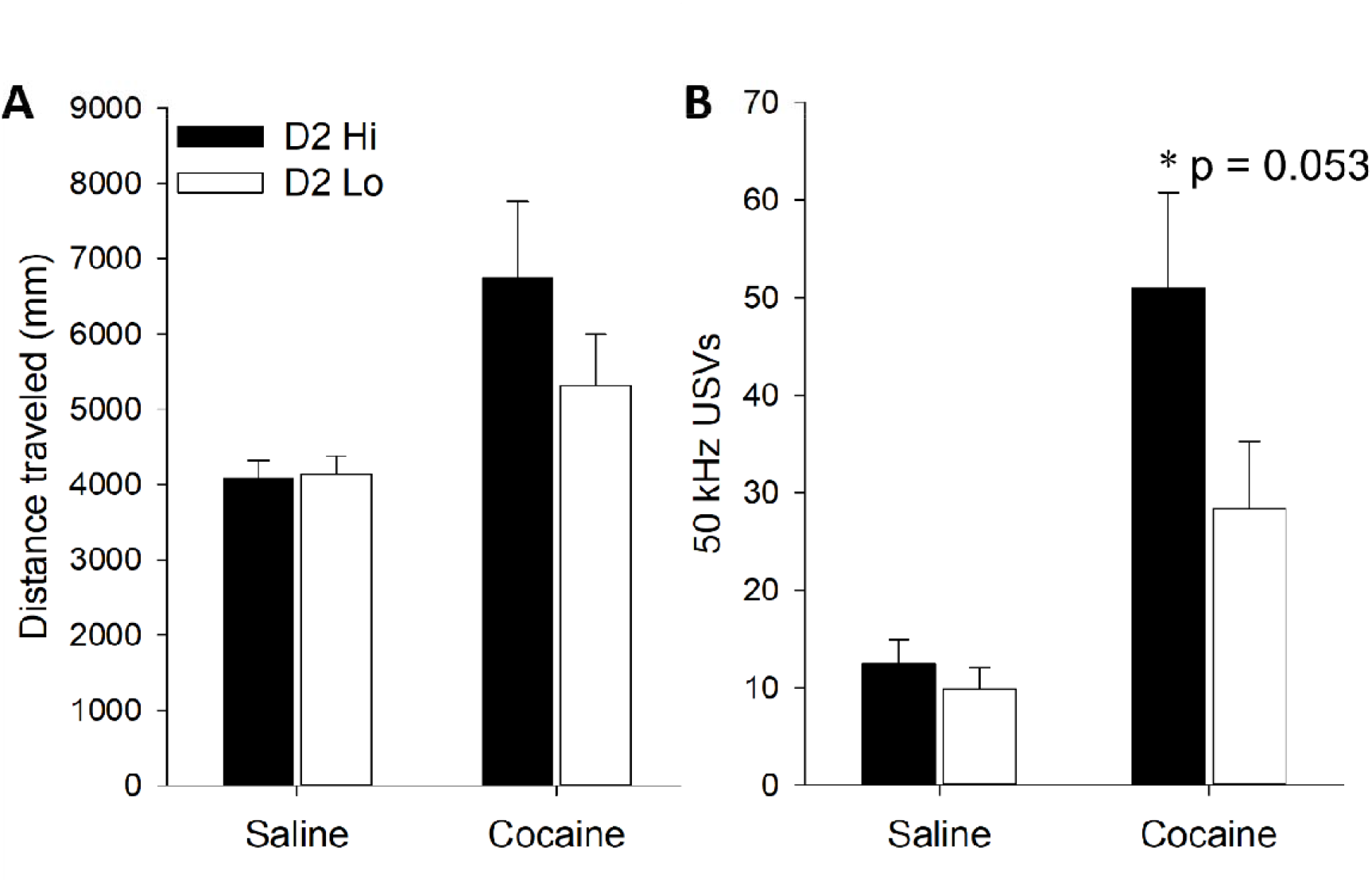
Locomotion (A) and USV production (B) in D2 Hi and D2 Lo rats following saline and cocaine treatment. Data are represented as the mean (+/-SEM). Asterisks (^*^) denote a trend towards a significant D2 sensitivity x Drug interaction (p = 0.053).

### Locomotor response to the D2 receptor agonist quinpirole and cocaine-induced behaviors in sign- and goal-trackers

As described above, D2 receptor sensitivity was related to PavCA behavior. In order to determine the converse, whether PavCA classification was related to sensitivity to the D2 receptor agonist and cocaine-induced behaviors, we classified individual rats as sign-trackers (ST) and goal-trackers (GT) based on their average PavCA index scores on days 4 and 5. Individuals with an average index score above 0.5 were classified as ST (n = 14) and below -0.5 were classified as GT (n = 14). One goal-tracker with extreme locomotor scores was removed as a statistical outlier.

STs and GTs differed in their response to quinpirole, as indicated by a Dose x Time x Phenotype interaction [F (33,858) = 1.83, p < 0.01], and post-hoc analyses indicated that STs were more sensitive to the 1.0 mg/kg dose during the peak locomotor response at 45 min (Fig. 4A; p < 0.05). STs were also more active during the first 5 minutes after saline injection (p < 0.01). There were no ST/GT differences in cocaine-induced locomotion (Fig. 4B), although STs produced more cocaine-induced USVs compared to GT [Fig. 4C; F (1,25) = 15.21, p < 0.001 for the Drug x Phenotype interaction]. This is a replication of our previous findings (Meyer et al., 2012b; Meyer & Tripi, 2018; Tripi et al., 2017) demonstrating a robust relationship between sign-tracking and cocaine-induced USVs.

**Fig. 4.**
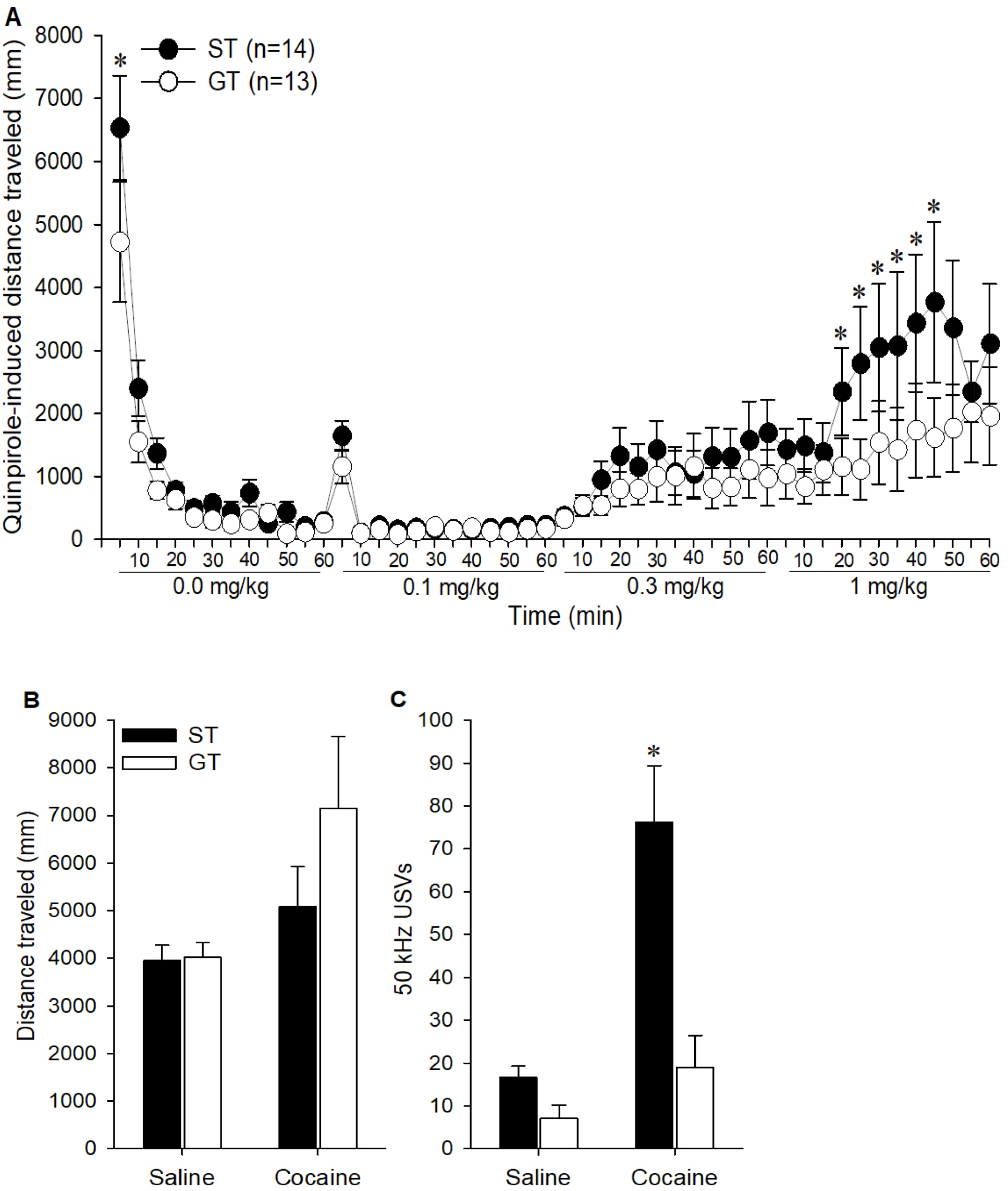
Rats characterized as sign-trackers (STs; n = 14) and goal-trackers (GTs; n=13) were compared in quinpirole-induced locomotion (A), cocaine-induced locomotion (B) and 50 kHz USV production (C). Data are represented as the mean (+/-SEM). Asterisks (^*^) denotes group differences (p < 0.05).

### Locomotor Response to the D1 receptor agonist SKF 82958 in sign- and goal-trackers

Then, only STs and GTs were injected with the D1 receptor agonist SKF 82958 to determine if they also differed in D1 receptor sensitivity. STs and GTs differed in their response to SKF 82958, as indicated by a 3-way interaction of Dose x Time x Phenotype [F (18,450) =1.67, p < 0.05], and post-hoc analyses indicated that STs were more sensitive to 1 mg/kg SKF 82958 dose during 35-50 min post-injection (Fig 5; p < 0.01). However, when rats were divided into high or low SKF 82958 responders using the median split method, high D1 (D1 Hi) and low D1 (D1 Lo) responders did not differ in cocaine-induced locomotion or USVs (data not shown).

**Fig. 5.**
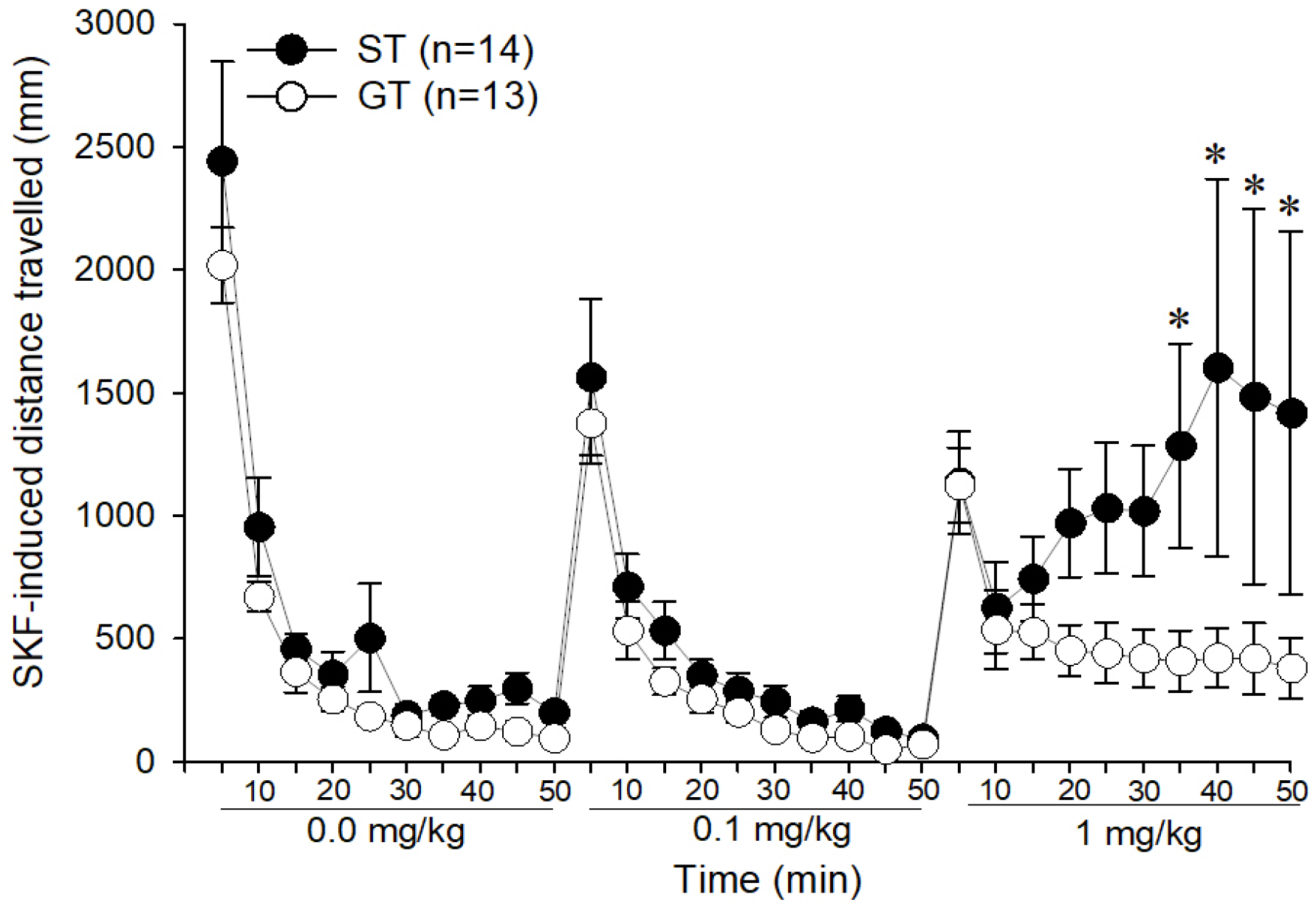
Locomotor response during D1 agonist dose-response test in sign- and goal-trackers (ST, GT; respectively). Data are represented as the mean (+/-SEM). Asterisks (^*^) denotes time-points of group differences (p < 0.05).

### Role of both D1 and D2 dopamine receptor sensitivity in PavCA and cocaine measures

Based on the pattern of results, we sought to determine whether sensitivity to both D1 and D2 receptor agonists was predictive of the PavCA and cocaine behaviors measured. Individuals tested with both quinpirole and SKF 82958 were grouped based on their Hi/Lo assignment for both agonists resulting in 4 groups: D1 Hi/D2 Hi (n = 6), D1 Hi/D2 Lo (n =7), D1 Lo/D2 Hi (n = 8) and D1 Lo/D2 Lo (n = 6). Using D1 sensitivity and D2 sensitivity as separate independent variables, we found that D2 Hi rats, regardless of D1 sensitivity, had more lever contacts than D2 Lo rats [main effect of D2 sensitivity: F (1,23) = 6.28, p < 0.05]. For food cup entries, both D1 and D2 sensitivity influenced this measure of goal-tracking [Day x D1 x D2 interaction: [F (4,92) = 4.13, 0.01; main effect of D2: F (1,23) = 6.06, p < 0.05]. Similarly, for the PavCA index score, both D1 and D2 sensitivity interacted to promote higher index scores, as indicated a 3-way interaction of Day x D1 x D2 [F (4,93) = 3.14, p < 0.05] and a main effect of D2 [F (1,23) = 8.65, p < 0.01). Post-hoc analyses indicated that Hi D2 rats, regardless of D1 sensitivity, had higher lever contacts, lower food-cup entries, and higher PavCA index scores than Lo D2 rats. Further, rats sensitive to both agonists (Hi/Hi) had higher PavCA index scores on days 4 and 5 (Fig 6A-C). This indicates that sensitivity to both D1 and D2 receptor drugs together is the best predictor of elevated levels of PavCA index scores.

**Fig. 6.**
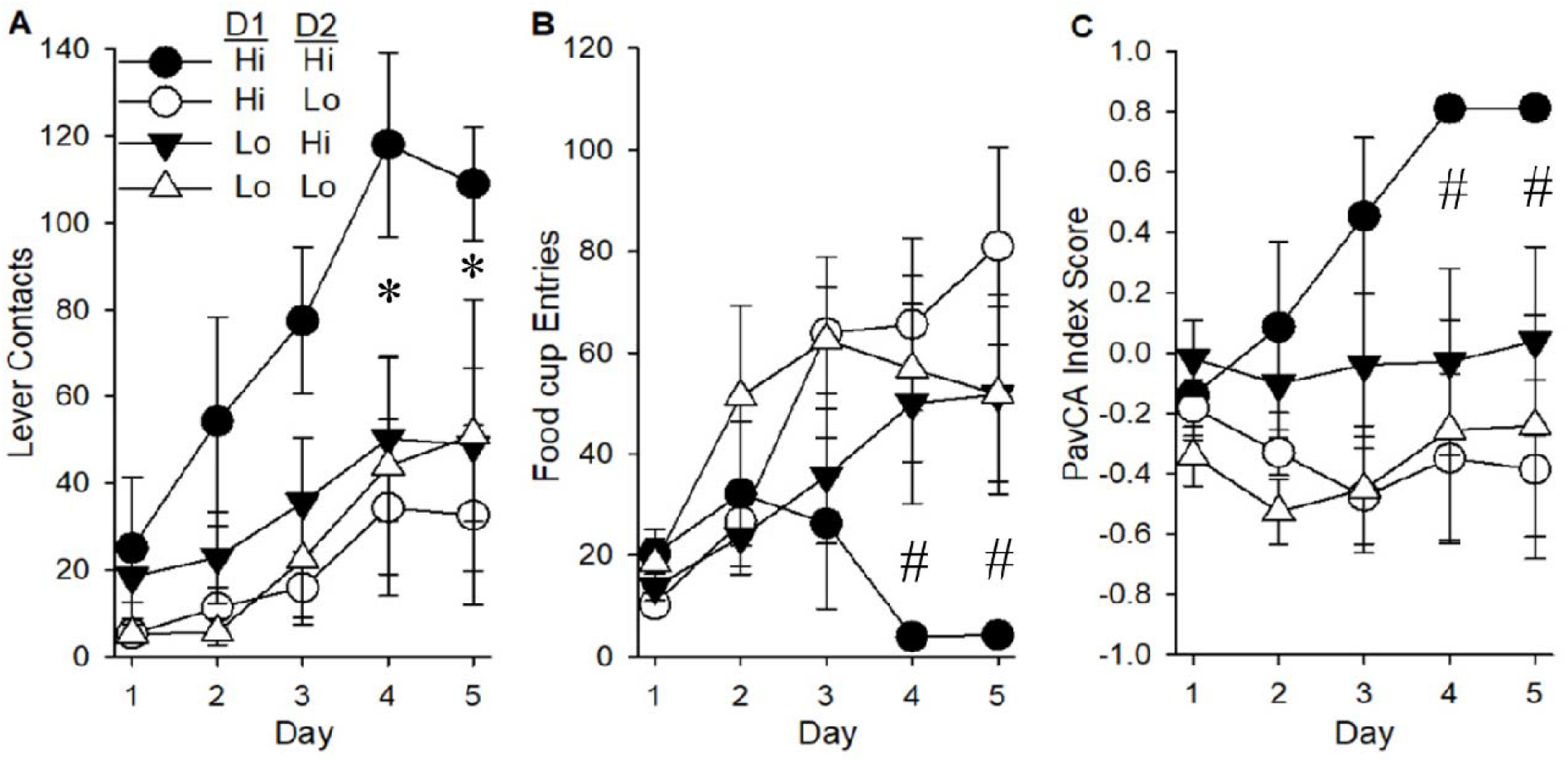
Pavlovian conditioned approach measures in rats characterized based on sensitivity to both D1 and D2 agonists. For the four resulting groups, D1 Hi/D2 Hi (n=6), D1 Hi/D2 Lo (n=7), D1 Lo/D2 Hi (n=8) and D1 Lo/D2 Lo (n=6), mean number of lever contacts (A) and food-cup entries (B) are shown. Mean PavCA index scores (C) were determined using the formula presented in “Methods and materials.” Data are represented as the mean (+/-SEM). Asterisks (^*^) denote 2-way interaction with Hi D2 groups having more lever contacts than Lo D2 groups (p < 0.05). Pound (#) denotes 3-way interaction with Hi/Hi group having lower food-cup entries and higher PavCA values on days 4 and 5.

Differences in locomotor and USV response to cocaine were also assessed in these four sensitivity groups (Fig 7A-B). There were no significant differences in cocaine-induced locomotion, but for cocaine-induced USVs there was a significant Drug x D1 x D2 interaction on USV production [F (1,23) = 5.91, p < 0.05]. Post-hoc analyses indicated that cocaine induced USVs by the Hi/Hi group were significantly higher than all other groups. Thus, when considered together, sensitivity to both dopamine D1 and D2 receptors is strongly associated with individual differences in PavCA behaviors and cocaine induced USVs.

**Fig. 7.**
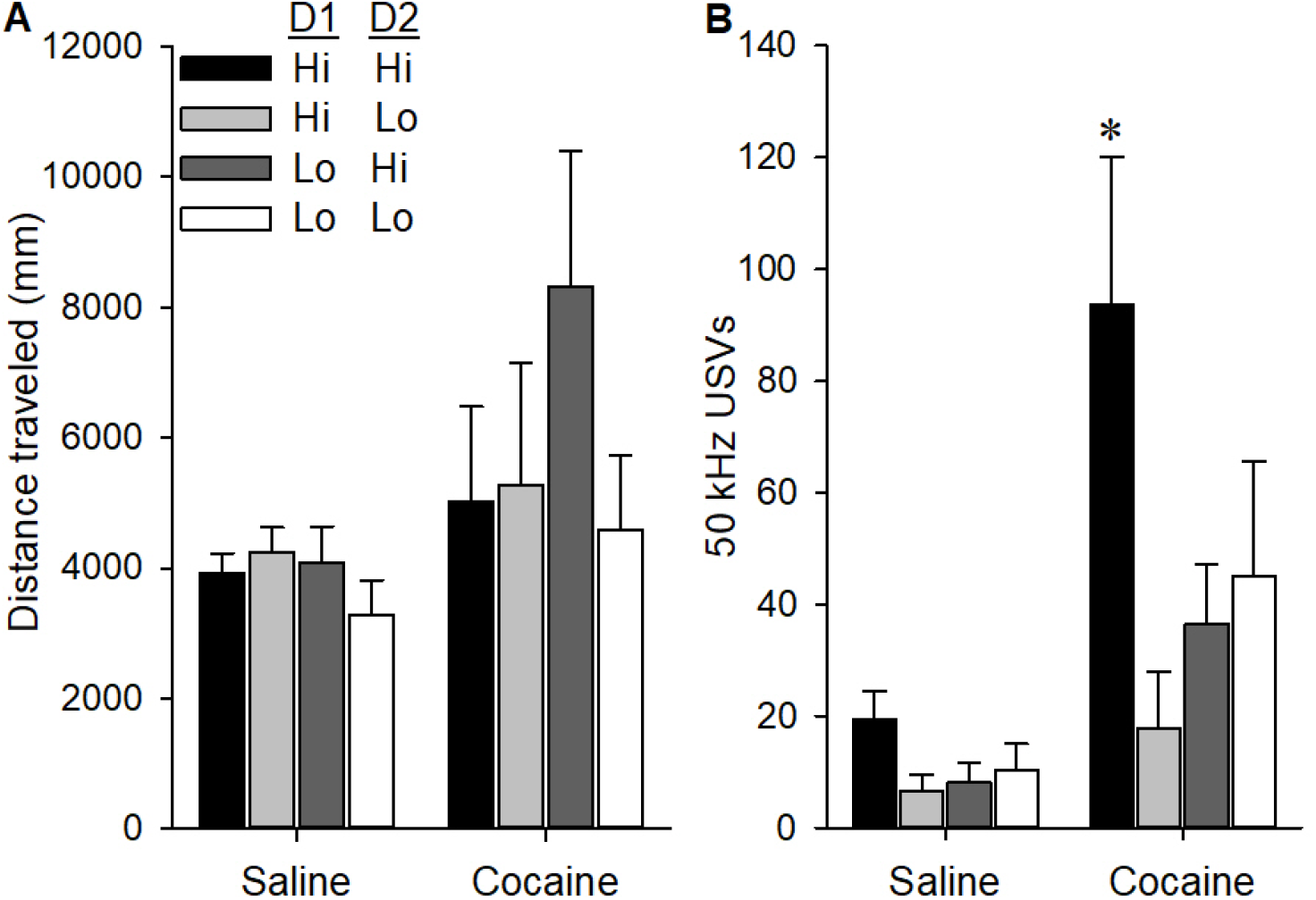
Locomotion (A) and USV production (B) in four sensitivity groups following saline and cocaine treatment. Data are represented as the mean (+/-SEM). Asterisks (^*^) denotes 3-way interaction with Hi/Hi group differing from all other groups (p < 0.05).

## Discussion

This study examined how dopamine D1 and D2 receptor sensitivity relates to reward cue-reactivity, and the unconditioned effects of cocaine in male rats. We found that rats with high quinpirole sensitivity (D2 Hi) were more likely to engage in sign-tracking compared to those with low sensitivity (D2 Lo). Cocaine increased locomotor activity in all rats, but there was a trend for greater cocaine-induced 50 kHz USVs in D2 Hi rats. Sign-trackers also showed more 50-kHz USVs than goal-trackers, but there was no difference in cocaine-induced locomotion. This indicates a robust association between sign-tracking and USVs. We also measured sensitivity to the D1 receptor agonist SKF 82958 and found that sign-trackers also had greater sensitivity to this drug. Additionally, subjects with high sensitivity to both D1 and D2 receptor drugs were most likely to sign-track and produce 50 kHz USVs. Together, these findings show that heightened D1 and D2 receptor sensitivity may lead to a predisposition to attribute incentive salience to reward cues, and to an enhanced affective response to cocaine.

### Predicting cue-reactivity based on D1 and D2 receptor sensitivity

Dopamine receptors play complex roles in learning and motivation (e.g. Baik, 2013; Larson et al., 2019; Nelson & Killcross, 2013; Rauhut et al., 2010; Roughley et al., 2021; Thibeault et al., 2019). Our results demonstrate that dopamine D2 receptor sensitivity predicts the tendency to attribute incentive to reward cues as measured by sign-tracking, while the lack of an effect on goal-tracking suggests that this was not due to an overall effect in learning. Similarly, high D1 and D2 receptor sensitivity was related to increased sign-tracking responses, and this was not due to enhanced learning overall, as indicated by a concomitant *decrease* in goal tracking. This evidence shows the involvement of both dopamine receptor subtypes in reward cue-reactivity, and aligns with earlier studies measuring dopamine receptor expression and pharmacological blockade of dopamine receptors (Saunders & Robinson, 2012). Specifically, Flagel et al. (2007) showed that D1 receptor mRNA was larger in sign-trackers after the first day of conditioning, whereas expression of other dopaminergic genes, including the D2 receptor, tyrosine hydroxylase, and dopamine transporter were increased in goal-trackers. This latter finding is somewhat incongruent with our finding of higher D2 receptor sensitivity in sign-trackers. However, these previous findings do not distinguish between pre- and post-synaptic receptor expression and do not measure potential differences in signaling pathways, which likely mediate the differences in sensitivity observed in our current study.

Studies with the nonspecific dopamine receptor antagonist α-flupenthixol and the D2 antagonist eticlopride show that dopamine receptor stimulation is required for the *expression* of both sign- and goal-tracking, but only the *acquisition* of sign-tracking is affected by these drugs (Flagel et al., 2011; Lopez et al., 2015; Roughley & Killcross, 2019). Drugs that antagonize D2 receptors, including olanzapine, haloperidol, and eticlopride, similarly blocked the acquisition of sign-tracking to a greater degree than goal-tracking, while the D1-receptor specific antagonist SCH39166 affected the acquisition of both conditioned responses (Danna & Elmer, 2010; Flagel et al., 2011; Roughley & Killcross, 2019). It is not clear why this drug affected goal-tracking but flupenthixol did not, but it is worth noting that SCH39166 appeared to affect sign-tracking more strongly than goal-tracking in (Roughley & Killcross, 2019). Lopez et al. (2015) also found that quinpirole selectively impaired the expression of sign-tracking, while Fraser et al. (2016) demonstrated that combined D2 and D3 receptor activation with 7-OH-DPAT and pramipexole attenuated expression of both sign- and goal-tracking responses. In this latter study, the authors argued that these drugs elicited this effect by disrupting endogenous signaling through these receptors. Our data are consistent with this explanation, in particular because the agonists used in our study were not given during the PavCA paradigm. In fact, our results extend prior work by demonstrating that individual differences in locomotor sensitivity to quinpirole and SKF 82958, inasmuch as this reflects variation in endogenous dopamine receptor signaling, are predictive of sign-tracking behavior. While falling short of demonstrating a causal role, this further emphasizes dopamine’s involvement in the attribution of incentive salience to reward-paired cues, and point a role for dopamine receptor sensitivity in determining the degree of cue-reactivity.

Neurocircuitry studies of the mesolimbic dopamine system support this. Nodes of this circuit, including the VTA, nucleus accumbens, and ventral pallidum, are more active during sign-tracking behavior than goal-tracking (Ahrens et al., 2016; Flagel et al., 2011; Gillis & Morrison, 2019; Saunders et al., 2018) and inhibition of dopamine neurons in the VTA blunted tendency to sign-track (Herring et al., 2025; Iglesias et al., 2023). Together with the pharmacological studies reviewed above, this indicates that individual differences in incentive salience attribution are likely related to both pre- and post-synaptic variations in the mesolimbic dopamine system.

### Dissociating cocaine induced-USVs and cocaine-induced locomotion

In our previous studies examining the relationship between cocaine-induced 50 kHz USVs and sign-tracking (Meyer et al., 2012b; Tripi et al., 2017), we found that sign-tracking rats emitted more cocaine-induced 50 kHz USVs than goal-trackers, which is consistent with previous finding demonstrating that 1) the dopamine system is altered in sign-trackers (Flagel et al., 2007, 2011), 2) sign-trackers are more sensitive to the incentive motivational effects of cocaine in several paradigms (Pitchers et al., 2017; Saunders & Robinson, 2010; Tunstall & Kearns, 2015), and 3) USVs may reflect the affective states associated with psychostimulant administration (Barker et al., 2014; Mahler et al., 2013). We also demonstrated the locomotor response to cocaine was *not* related to sign-tracking, which was intriguing because, since cocaine’s unconditioned effects are thought to be primarily mediated by its ability to increase extracellular dopamine, one might expect that cocaine-induced USVs and cocaine-induced locomotion would be related.

Therefore, in this study, we wondered whether this dissociation may be due to differential sensitivity to D1 and D2 dopamine receptor stimulation among sign and goal-trackers. This appears to be the case. Sign-tracking was positively associated with cocaine-induced 50 kHz USVs as we had previously found, and also positively associated with locomotor sensitivity to D1 and D2 receptor stimulation. However, sign-tracking was again not associated with cocaine-induced locomotion. Further, rats that were sensitive to the locomotor effects of both D1 and D2 receptor stimulation were more likely to emit cocaine-induced USVs, but were *not* more likely to show more cocaine-induced locomotion.

Findings from this study and previous studies examining D1 and D2 receptor sensitivity shed some light on why cocaine induced USVs and locomotion are dissociable. D1 and D2 receptor sensitivity were related to cocaine-induced USVs but not cocaine-induced locomotion. Like others, (Scardochio & Clarke, 2013) we did not observe any increases in 50 kHz vocalizations induced by D1 and D2 receptor agonists, even though both quinpirole and SKF 82958 increased locomotion. There are several possible explanations for this. First, simultaneous activation of D1 and D2 receptors may be required to elicit USVs, but not locomotion. Second, cocaine may cause vocalizations via a non-dopaminergic mechanism, require simultaneous activation of both receptors or produce dynamic (phasic vs tonic) changes in dopamine that differentially affect USVs. Cocaine and other psychostimulants such as amphetamine increase norepinephrine in multiple brain areas through their actions on the norepinephrine transporter (Berridge & Stalnaker, 2002; Florin et al., 1994). These noradrenergic actions appear to be an underlying factor for amphetamine-induced USVs (Wright et al., 2013), and are likely similarly involved in cocaine-induced USVs. Complicating this explanation, however, is the demonstration that these adrenergic actions are involved in psychostimulant-induced locomotion as well (Darracq et al., 1998; Drouin et al., 2002). Thus, it is still difficult to explain why D1 and D2 sensitivity is associated with cocaine-induced USVs and not cocaine-induced locomotion. Given that drug-induced locomotion is governed by multiple neurochemical substrates (Holstein et al., 2005; Meyer & Phillips, 2003; Phillips & Shent, 1996; Wise & Bozarth, 1987).

Another explanation relates to cocaine’s dynamic (phasic vs tonic) changes in dopamine signaling that might differentially affect USVs and locomotion. Psychostimulants enhance phasic, sub-second dopamine transmission (Aragona et al., 2008; Cheer et al., 2007; P. E. M. Phillips et al., 2003; Ramsson et al., 2011), as well as long term dopamine increases (Di Chiara, 2002; Meyer et al., 2009). As such, cocaine-induced USVs may be due to its effects on phasic dopamine neurotransmission, whereas cocaine-induced locomotion may be due to its effects on tonic dopamine (and noradrenergic) transmission. Supporting this, optogenetic stimulation of dopaminergic neurons in the nucleus accumbens (NAcc) increased 50-kHz USVs, and these USVs were similar to psychostimulant induced USVs (Scardochio et al., 2015). In another study, Scardochio and Clarke (2013) found that D1 and D2 receptor activation inhibited USVs induced by amphetamine, suggesting a blocking or masking of the postsynaptic impact of this phasic transmission, which would explain why these drugs do not induce USVs, but still increase locomotion due their tonic release of dopamine (Freed & Yamamoto, 1985) and subsequent stimulation of these receptors. While speculative, our data are consistent with this argument as well. Unfortunately, it is not known whether phasic increases dopamine transmission are related to individual differences in dopamine receptor sensitivity than tonic increases.

### Dopamine receptor sensitivity and sensitivity to cocaine

In this study, high sensitivity to *both* D1 and D2 receptor stimulation was related to increased cocaine-induced locomotion and cocaine-induced 50 kHz USVs, but unlike Merritt and Bachtell (2013), we did not see a robust relationship between D2 receptor sensitivity alone and the locomotor response to cocaine. This may be due to the slightly lower dose in our study (10 mg/kg), because Merritt and Bachtell found more robust effects with 15 mg/kg cocaine. This could also be due to parametric differences. For example, our locomotor chambers are smaller, which may have prevented observation of robust group differences. If this is the case, then it seems likely that the individual differences in sign-tracking during PavCA, as well as the locomotor, USV, and motivation responses to cocaine are reflective of an underlying difference in sensitivity to D2 (and D1) receptor activation. Supporting this, other studies suggest that sign-trackers are more sensitive to the incentive motivational effects of cocaine in several paradigms. Saunders and Robinson (2010) showed that cocaine cues stimulated drug taking, and reinstated string drug-seeking (post extinction) specifically in rats that sign-tracked during PavCA. Tunstall and Kearns (2015) also backed this by demonstrating that compared to GTs, STs were more likely to choose a cocaine reward over food reward. There is additional evidence to suggest that cue-induced cocaine seeking is stronger in STs (Saunders et al., 2013), who have higher motivation for cocaine (Saunders & Robinson, 2011), and show greater locomotor sensitization to cocaine (Flagel et al., 2007). Finally, incentive value attribution is associated with other addiction-related traits, including impulsivity (King et al., 2016; Lovic et al., 2011; Yates & Bardo, 2017) and suboptimal choice (Chow et al., 2017; López et al., 2018; but see Orduña & Alba, 2020). Thus, the combination of findings from this study and others suggests that assigning incentive value to reward cues could be considered a phenotypical marker of addiction-like traits, and may be mediated by sensitivity to D1 and D2 receptor stimulation.

## Conclusion

In conclusion, there is an extensive body of research connecting changes in the mesolimbic dopaminergic system to both the use of psychostimulants and subsequent effects, with sensitivity and expression of D2 receptors considered to be associated with vulnerability and response to psychostimulants (Moreira & Dalley, 2015; Salery et al., 2020). Merritt and Bachtell (2013) backed this by reporting higher cocaine locomotor activity and place preference for rats with high D2 receptor sensitivity. Our findings extend this work by showing that individual variation in D1 and D2 receptor sensitivity not only predicts sensitivity to cocaine, but may also underlie stable, trait-like differences in cue-reactivity and susceptibility to addiction-related behaviors.

## Acknowledgement

The authors would like to acknowledge Dr. Micheal Dent of the Department of Psychology (University at Buffalo) for contributing to this project. Dr. Dent provided the equipment for USV recording, and was consulted on USV data analyses. This work was supported by grants from the Office of the Director, National Institutes of Health, the National Institute on Alcohol Abuse and Alcoholism, and the National Institute of Drug Abuse U01DA060669, P50DA037844, R01AA024112, T32AA007583.

